# More than meets the Kappa for Antibody Superantigen Protein L (PpL)

**DOI:** 10.1101/2021.11.26.470168

**Authors:** Wei-Li Ling, Joshua Yi Yeo, Yuen-Ling Ng, Anil Wipat, Samuel Ken-En Gan

## Abstract

Immunoglobulin superantigens play an important role in the affinity purification of antibodies and underlie the microbiota-immune axis at mucosal areas Focussing on the *Staphylococcal* Protein A (SpA), *Streptococcal* Protein G (SpG), and the *Finegoldia* Protein L (PpL) that were previously thought to bind to only specific regions of human antibodies, a systematic and holistic analysis of the antibody regions using 63 antibody permutations involving six Vκ and seven VH region IgG1 revealed showed novel PpL-antibody interactions. While SpA and SpG showed relatively consistent interactions with the antibodies, our findings showed PpL binding to certain VH-Vκ2, 5 and 6 interactions had contribution by other antibody regions. The findings of this have implications on PpL-based affinity antibody purifications and antibody design as well as provides novel insights to PpL-based microbiota-immune axis effects.

## 1 Introduction

B-cell superantigens bind antibodies or immunoglobulins (Ig) to hyperstimulate populations of B-cells independent of T-cells and have been used widely used for antibody affinity purification (Deacy et al., 2021). Superantigens are predominantly produced by microorganisms as a defence mechanism to escape from the host immune system (Spaulding et al., 2013). Notably there are three widely-used antibody superantigens also known as immunoglobulin binding proteins (IBP): Protein G (SpG) which binds the heavy chain constant region of the IgG subtypes (IgG1-4) and is produced the by groups C and G of *Streptococcal* bacteria (Sjöbring et al., 1991); Protein A (SpA) produced by *Staphylococcus aureus* which also binds to the heavy chain constant region of IgG1, 2, and 4 and also the variable heavy (VH) 3 framework (VH3) (GROV et al., 1964;Sasso et al., 1991;Su et al., 2021); and Protein L (PpL) produced by *Finegoldia magna* (previously known as *Peptostreptococcus magnus*) which binds to the variable light kappa κ (Vκ) chain families 1,3,4 (Nilson et al., 1992) at the framework (FWR) 1 region with influence by the other regions (Su et al., 2017).

When bound to the antibodies, these superantigens can reduce the binding of the antibodies to their antigen (Ling et al., 2021), possibly reducing avidity through steric hindrances as in the case of IgM (Samsudin et al., 2020), cause unwanted activation (Su et al., 2021) with downstream effects depending on their isotype (discussed in (Ling et al., 2020a;Gan, 2021)).

With both IgG and Vκ as the predominant isotypes in humans (Haraldsson et al., 1991;Janeway et al., 2001), superantigen proteins A, G, and L are likely to underlie significant microbiota-immune axis interactions especially at colonization of mucosal areas. Nonetheless, the problem extends to antibody purification process at which these superantigens are often used. Considering that most therapeutic antibodies are of the IgG and κ isotypes, unwanted interactions of such superantigens produced by commensals at the natural colonization sites can influence the microbiota-immune axis.

Given the widespread implications of these three superantigens, a holistic (Phua et al., 2019;Ling et al., 2020a) and systematic antibody-superantigen investigation using 63 of our previously engineered antibodies (Ling et al., 2018;Lua et al., 2018;Ling et al., 2020b) was performed involving six Vκ and seven VH IgGs finding novel interactions for PpL but not for SpA and SpG.

## 2 Materials and Methods

### 2.1 Recombinant Antibody Production

All Trastuzumab and Pertuzumab VH and Vκ sequences used were described previously (Ling et al., 2018;Ling et al., 2020b). Briefly, the genes were sub-cloned into pTT5 vector (Youbio, Cat: VT2202) using restriction enzyme sites, as previously performed (Su et al., 2017;Ling et al., 2018;Lua et al., 2018;Lua et al., 2019a;Ling et al., 2020b). The plasmids were transformed into competent *E. coli* (DH5α) strains (Chan et al., 2013) followed by plasmid extraction (Biobasic Pte Ltd, Cat: BS614).

Transfection, production, and purification and were performed as described previously (Ling et al., 2018;Ling et al., 2020b).

### 2.2 Binding Affinity Quantification

Measurements of superantigen ka and kd using the OctetRed® were performed using PpL (Sartorius, Cat: 18-5185), SpA (Sartorius, Cat: 18-5012), and SpG (Sartorius, Cat: 18-18-5083) biosensor to Pertuzumab and Trastuzumab IgG1 variants in solution. The program and steps used were as previously described (Su et al., 2017;Ling et al., 2018;Lua et al., 2018;Su et al., 2018;Lua et al., 2019b;Ling et al., 2020b;Su et al., 2021).

## 3 Results

### 3.1 Bio-Layer Interferometry (BLI) measurement of recombinant IgG1 variants to Protein A (SpA)

To examine the potential holistic effect of Vκ1-6 pairing with VH1-7 on antibody interactions with SpA, recombinant Pertuzumab and Trastuzumab IgG1 variants of the various pairings were studied. It should be noted that SpA is known to bind to the CH2 and CH3 of the heavy chain constant (CH).

From Figure 1, the Pertuzumab and Trastuzumab IgG1 variants showed measurements bound to SpA with the equilibrium dissociation constant (KD) at 0.41 – 1.90 × 10^−9^ M and 0.25 – 2.30 × 10^−9^ M, respectively.

**Figure 1.**
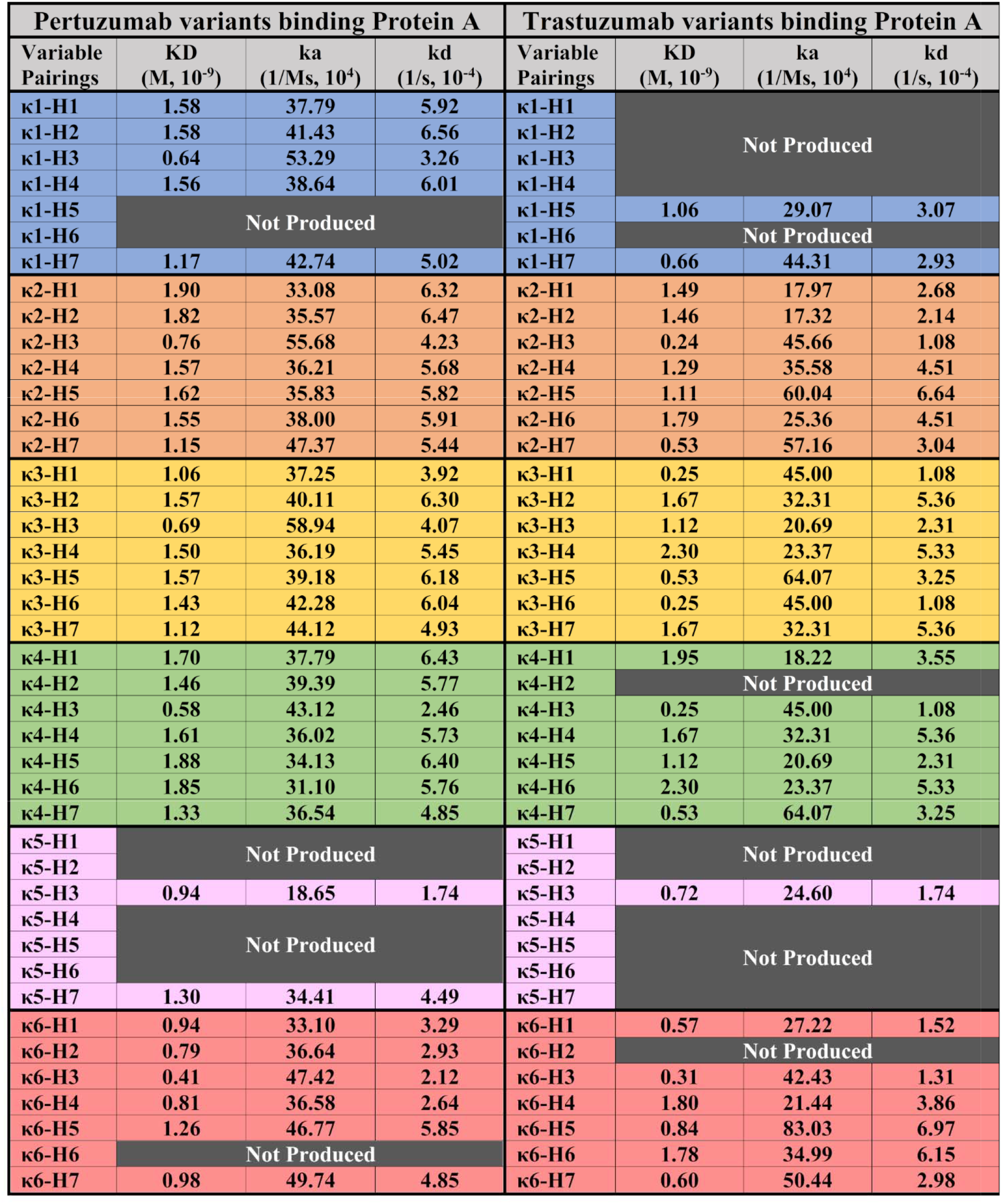
BLI measurements (KD, ka and kd) of Pertuzumab and Trastuzumab Vκ1-6 and VH1-7 permutation binding to immobilized SpA biosensor. “Not Produced” denotes that there was insufficient antibody production for the variant despite numerous large-scale transfections. All readings were obtained from at least three antibody concentrations. The readings were the average of independent triplicates.

For both Pertuzumab and Trastuzumab variants binding to SpA, VH3 was noticed to have a slightly lower, albeit insignificant KD difference at the average of ~0.57 × 10^−9^ M and ~0.53 × 10^−9^ M, respectively, when compared to the other VH families paired with the same Vκ family. This phenomenon is attributed to the higher ka and lower kd for the VH3 variant.

### 3.2 Bio-Layer Interferometry (BLI) measurement of recombinant IgG1 variants to Protein G (SpG)

Testing the 63 recombinant Pertuzumab and Trastuzumab IgG1 variants with SpG which binds to CH2 and CH3 region, we found high consistency of the interactions between the two Pertuzumab and Trastuzumab IgG1 variants. Apart from showing similar KDs to the SpA, albeit with narrower KD ranges of 0.23 – 0.87 × 10^−9^ M and 0.23 – 1.78 × 10^−9^ M, respectively. There is a trend of Pertuzumab variants binding SpG better than the Trastuzumab counterparts. This slight difference, while unlikely significant, hints of CDR effects given that the variants differed only at a few residues in the CDRs.

### 3.3 Bio-Layer Interferometry (BLI) measurement of recombinant IgG1 variants binding to Protein L (PpL)

From the total 63 permutations IgG1 variants consisting of 34 Pertuzumab and 29 Trastuzumab permutations of the grafted Vκ1-6 and VH1-7 on PpL, our systematic and holistic investigation of IgG1s to PpL showed non-canonical results of interactions with other Vκ families and a contributory role of VH-FWR and complementarity-determining regions (CDRs) to the interaction.

As a control for expected superantigen interactions, the Pertuzumab IgG1s of Vκ1, 3 and 4 interacted with PpL with Vκ1 showing the lowest KD range (0.53 - 0.76 × 10^−9^ M) followed by Vκ3 (5.55 - 38.03 × 10^−9^ M) and Vκ4 (13.09 – 74.56 × 10^−9^ M). The Vκ1 findings were consistent with previous literature (Åkerström and Björck, 1989;Rodrigo et al., 2015;Su et al., 2017). The lower equilibrium dissociation constants (KDs) of the Vκ3 and 4 were due to the lower dissociation rates (kd) despite the higher association rates (ka) than Vκ1. This trend was also observed for the Trastuzumab IgG1s with its Vκ1 showing the lowest KD range (0.11 & 0.14 × 10^−9^ M) followed by Vκ3 (3.67 – 17.77 × 10^−9^ M) and 4 (7.22 – 14.18 × 10^−9^ M). It should be noted that Trastuzumab IgG1s showed lower and a narrower KD range than the Pertuzumab Vκ-VH equivalents suggesting effects from the CDRs which were what differed between the two sets of IgG1s.

Interestingly, certain Pertuzumab VHs paired with Vκ2, 5 and 6 exhibited interactions with PpL. Amongst these Vκ families, Pertuzumab variants Vκ5 (13.58 & 13.88 × 10^−9^ M), 6 (15.79 – 43.46 × 10^−9^ M) and certain Vκ2 permutations (VH2-4 and 7, 14.4 – 46.4 × 10^−9^ M) had KDs comparable to Vκ3 and 4 permutations (5.55 – 74.56 × 10^−9^ M). PertuzumabVκ2 paired with VH5 and 6 (0.93 and 0.72 × 10^−9^ M, respectively) had KDs comparable to Vκ1 (0.53 - 0.76 × 10^−9^ M) while Vκ2 paired with VH1 had the highest KD (poorest binding) of 148.67 × 10^−9^ M. There were also non-binding IgG1s of the Pertuzumab Vκ6 permutation with VH1-3 (Poor Response) despite measurable responses when paired with VH4, 5 and 7.

Interesting, the Pertuzumab trends were largely repeated in the Trastuzumab IgG1s where KDs of Vκ5 (1.75 × 10^−9^ M) and 6 (14.84 – 82.16 × 10^−9^ M) and Vκ2 paired with VH2 & 7 (78.63 & 9.79 × 10^−9^ M, respectively) had KD values comparable to Vκ3 and 4 (3.67 – 14.18 × 10^−9^ M). Trastuzumab Vκ2 paired with VH1 & 6 had the highest KD (poorest interaction) at 124.41 & 190.4 × 10^−9^ M, respectively. The non-binders were Trastuzumab Vκ2 paired with VH3 – 5 (Poor Response) rather than in Vκ6 family observed for Pertuzumab. These differences demonstrated a role of the CDRs and a significant contributory role of VH in PpL engagement.

## 4 Discussion

We set out to investigate the interactions of superantigens Protein A, G and L systematically and holistically with the various regions of IgG1 antibodies. By using CDRs of Pertuzumab and Trastuzumab grafted onto Vκ1-6 and VH1-7 FWRs and pairing them within the two antibody models, measurements to the antibody superantigens showed no major differences for SpA between Pertuzumab and Trastuzumab IgG1 variants (Figure 1). This was expected given that SpA bound IgG1s predominantly at the CH2-CH3 regions (Deisenhofer, 1981) with some contributions from the VH3 framework (Su et al., 2021) that is also observed here to a lesser extent where the Vκ chains paired with VH3 showed a slightly lower KD measurement compared to the rest of the variants. Yet, this difference is notably less pronounced compared to our previous work on IgEs (Su et al., 2021) with the same Vκs-VH where the VH3-CDR2 S58 residue had a more significant role in SpA binding for IgEs.

With respect to SpG interaction, no notable differences in KDs (Figure 2) were observed among the 63 IgG1 variants. As was with SpA, which shared an overlapping binding site on the CH2-CH3 region of IgGs (Kato et al., 1995), there was in fact a narrower range that we attributed to the lack of interference from the V-regions present for SpA. While SpG was previously reported (Choe et al., 2016) to bind to IgG1 better than SpA, this trend was more pronounced in our study.

**Figure 2.**
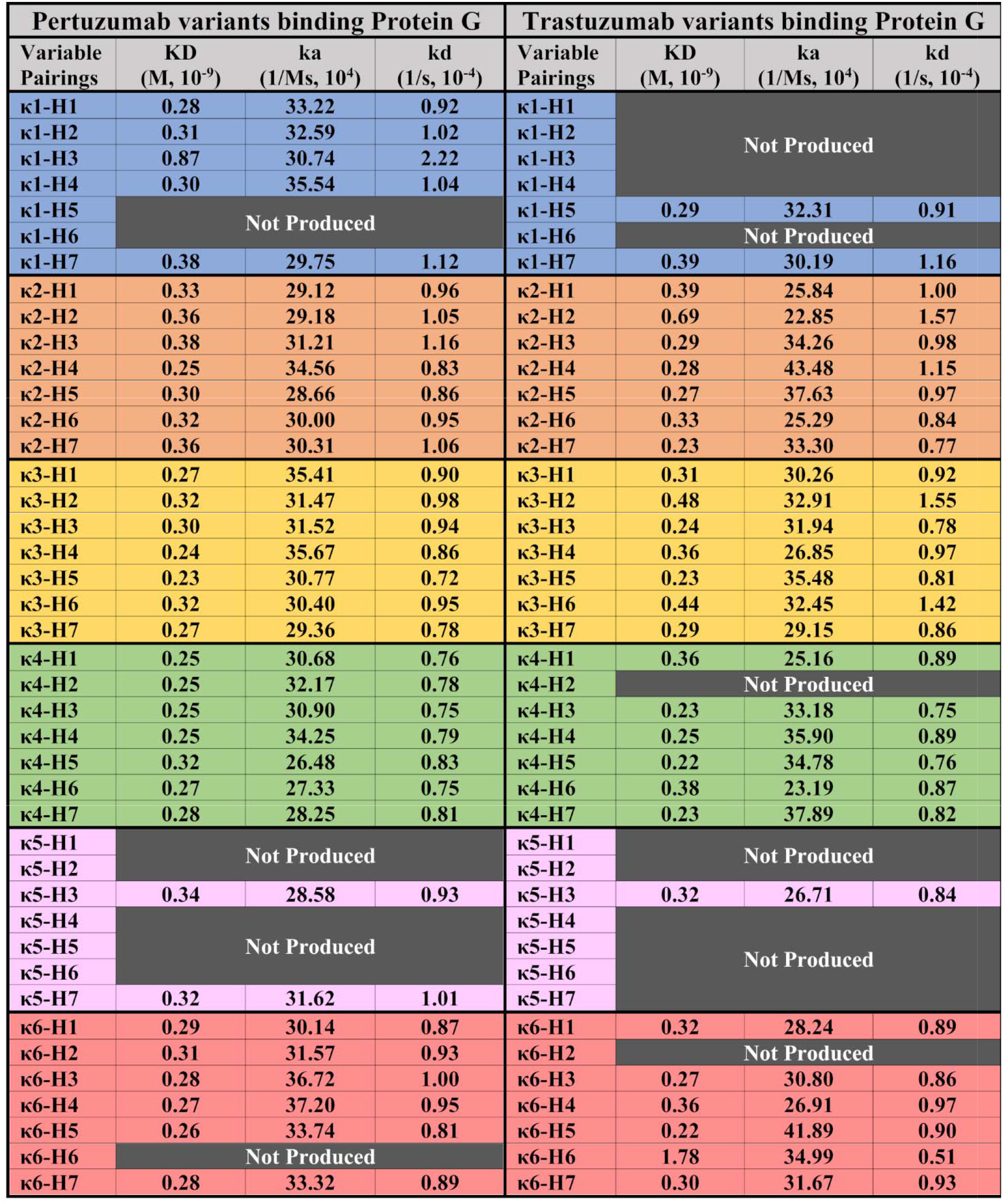
BLI measurements (KD, ka and kd) of Pertuzumab and Trastuzumab Vκ1-6 and VH1-7 permutation binding to immobilized SpG biosensor. “Not Produced” denotes that there was insufficient antibody production for the variants despite numerous large-scale transfections. All readings were obtained from at least three antibody concentrations. The readings were the average of independent triplicates.

Measurements of PpL interactions to Pertuzumab and Trastuzumab variants expectedly showed that VHs paired with Vκ1, 3 and 4 to have KDs as per previously reported (Nilson et al., 1992;Su et al., 2017). Surprisingly, we found non-canonical interaction of PpL with Vκ2 that were previously determined to not bind PpL (Nilson et al., 1992) while there is no report of Vκ5 & 6 at the time of writing. In our own work involving light chain productions alone, we also affirmed that these secreted Vκ2, 5 and 6 light chain dimers did not interact with PpL on the same BLI experiments (Supplementary Figure 1). Notable binders to PpL are: Pertuzumab Vκ2 – VH1-7; Vκ5 – VH3 & 7; Vκ6 – VH4, 5 & 6; Trastuzumab Vκ2 – VH1, 2, 5 & 6; Vκ5 – VH3; Vκ6 – VH1, 3-7 (Figure 3).

**Figure 3.**
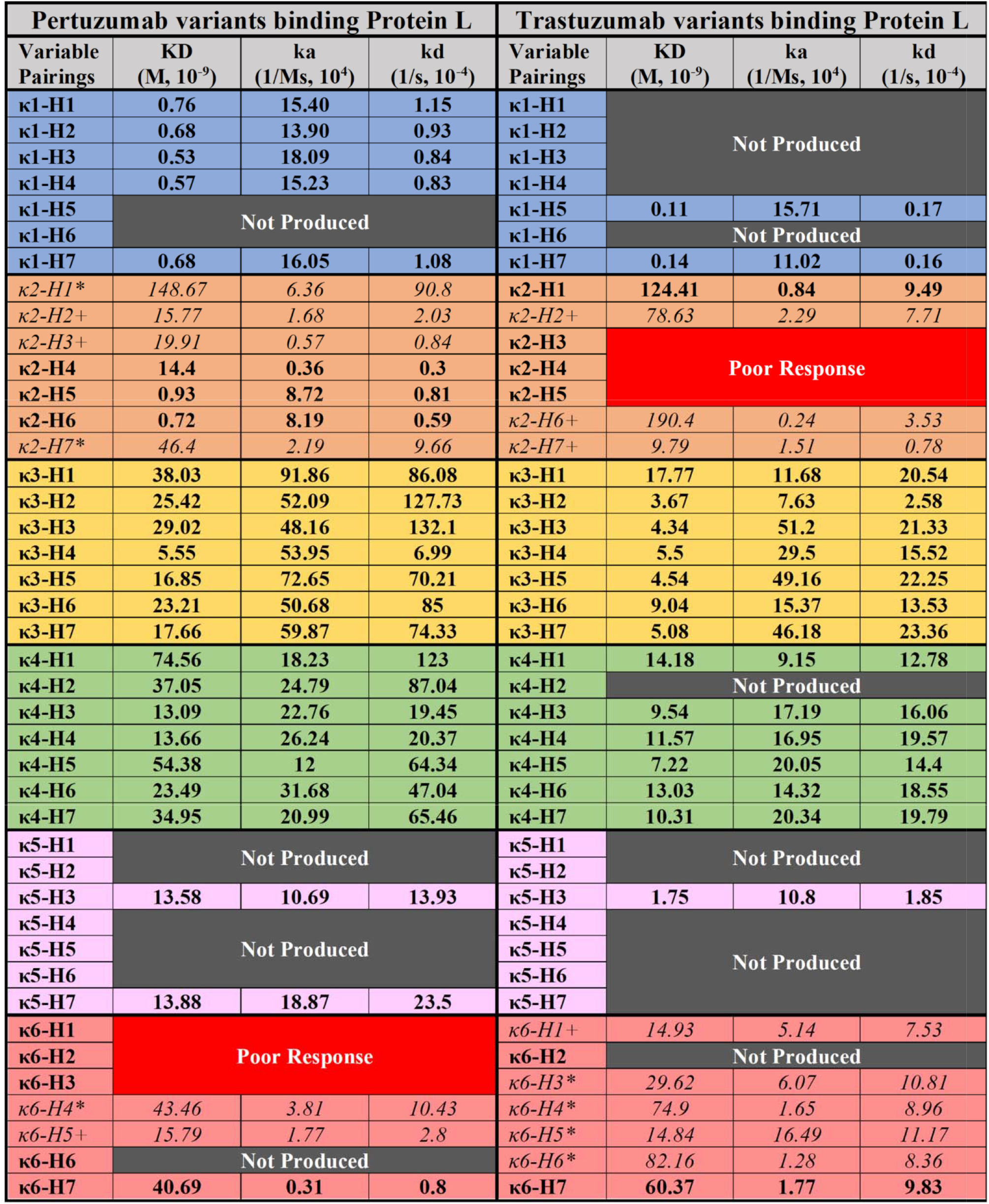
BLI measurements (KD, ka and kd) of Pertuzumab and Trastuzumab Vκ1-6 and VH1-7 permutation binding to immobilized SpG biosensor. “Not Produced” denotes that there was insufficient antibody production for the variants despite numerous large-scale transfections. “Poor response” indicates that the particular IgG1 pairing was unable to give response rates within the detection limit across all concentrations. * denotes readings that were derived from two IgG1 concentrations. + denotes represent readings generated derived from only one IgG1 concentration. All other readings were obtained from at least three concentrations. The readings were the average of independent triplicates.

Although the novel Vκ IgG1s bound PpL showed comparable KDs, it should be noted that the KDs were calculated from one (+ in Figure 3) or two (* in Figure 3) antibody concentrations, generally from the highest concentrations (100 nM and below) of the Ig variant. The notable exceptions were that of Pertuzumab Vκ2 – VH4-6, Vκ5 – VH3 & 7, Vκ6 – VH6, Trastuzumab Vκ2 – VH1, Vκ5 – VH3, Vκ6 – VH7 with KDs calculated from at least three concentrations. Interestingly, two variants: Pertuzumab Vκ2 – VH5 & 6 showed KDs comparable to Vκ1 – VHs values.

The unexpected IgG1 variants interacting with PpL suggested a combined VH-Vκ induced binding site to PpL that may be similar to the non-canonical binding of IgEs to Nickel (Su et al., 2021) in our previous work using the same V-regions. In fact, the IgG1s were validated with the expected interactions to SpA and SpG here, and also with the Fcγ2A and Her2 in our previous work (Ling et al., 2018). Given the lack of interactions between Vκ 2, 5 and 6 with PpL, and the lack of consistency between the Trastuzumab and Pertuzumab variants where for Pertuzumab, the non-binders exist for Pertuzumab Vκ6 – VH1-3 and Trastuzumab Vκ2 – VH3-5 (labelled as “Poor Response” pairs in Figure 3), the PpL interaction is certainly beyond V-region pairings alone.

With the differences between the Pertuzumab and Trastuzumab which share very similar V-regions, our findings further demonstrate the need for a design thinking (Ling et al., 2020a) approach involving holistic antibody investigations approach (Phua et al., 2019). Such an approach allowed detailed investigations for unexpected interactions between the antibodies with other proteins that can have notable immune effects, as was with our unexpected findings of IgAs binding to SpG (Ling et al., 2021). With relevance to the development of therapeutics where a personalized antibody approach may be beneficial to avoid unwanted side effects, such interactions may also be engineered in for purification purposes.

## 5 Conflict of Interest

*The authors declare that the research was conducted in the absence of any commercial or financial relationships that could be construed as a potential conflict of interest*.

## 6 Author Contributions

Conceptualization, W.L.L. and S.K-E.G.

Methodology, W.L.L. and S.K-E.G.

Investigation, W.L.L. and S.K-E.G

Validation, J.Y.Y.

Writing – Original Draft, W.L.L. and S.K-E.G

Writing – Review & Editing, W.L.L., Y.L.N. and S.K-E.G

Funding Acquisition, S.K-E.G.

Supervision, S.K-E.G., Y.L.N. and A.W.

## 7 Funding

This work was partially supported by the Joint Council Office, Agency for Science, Technology, and Research, Singapore under Grant number JCO1334i00050 and the National Research Foundation (NRF) Singapore grant to Experimental Drug Development Centre (EDDC).

## 8 Acknowledgments

This is a short text to acknowledge the contributions of specific colleagues, institutions, or agencies that aided the efforts of the authors.

## 10 Data Availability Statement

The datasets GENERATED/ANALYZED for this study is available upon request.

## Notes

### Competing Interest Statement

The authors have declared no competing interest.

