## Supplementary Figure 1 for "More than meets the Kappa for Antibody Superantigen Protein L (PpL)"

Supplementary Material

### 1 Supplementary Figures

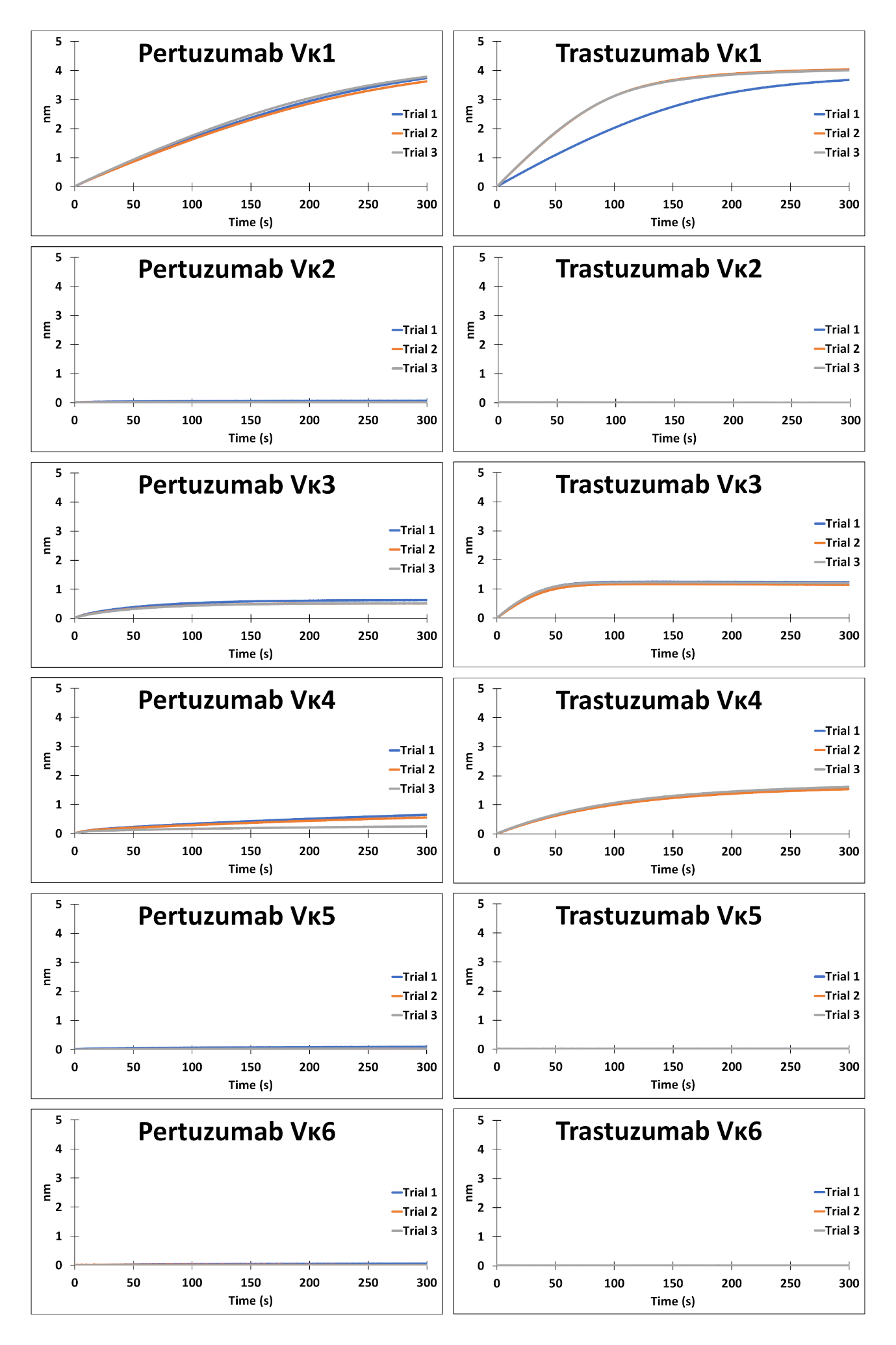

**Supplementary Figure 1.** BLI measurement of Pertuzumab and Trastuzumab Vκ1-6 binding to immobilized PpL biosensor.
